# Climate adaptation across space and time: lessons from oak populations

**DOI:** 10.64898/2026.08.06.743225

**Authors:** José A. Ramírez-Valiente, Joaquín Ortego, Antoine Kremer

## Abstract

Tree populations can respond to climate change through migration, phenotypic plasticity, or genetic evolution. Despite long generation times of forest tree species, recent studies suggest that their evolutionary responses may occur rapidly. Using oaks as a model system, we synthesize evidence from 88 common garden studies and from historical, retrospective and longitudinal approaches to explore how populations have adapted to climatic variability across different biomes, and assess the consistency and pace of evolutionary responses across spatial and temporal climatic gradients.

We found that approximately 61% of the studies exhibited significant differences among populations but climatic drivers and adaptive strategies differed among biomes. Temperature-related clines predominated in temperate regions, with populations from warmer origins showing longer growing seasons and higher growth potential. In seasonally dry biomes, aridity favored increased drought tolerance in Mediterranean populations but drought avoidance in tropical populations. Allochronic studies revealed genetic changes over decades to millenia in response to climate changes, with warming associated with increased growth and reduced specific leaf area in temperate oaks. Thus, spatial differentiation and temporal evolution were generally congruent in direction for most traits except for leaf unfolding, while short-term evolutionary rates exceeded long-term estimates by two to three orders of magnitude.

In summary, provenance trials can provide useful information on the direction of climate-driven evolution for some traits, but may underestimate its contemporary pace. More studies are needed to evaluate whether standing genetic variation of forest tree species is sufficient to track current climate change.

## 1. Introduction

Forests worldwide concentrate global environmental issues. Although they are expected to mitigate global greenhouse gas emissions by increasing the terrestrial carbon sink, their ability to meet these demands is, however, undermined by expanding tree mortalities caused by climate change [1–3]. The capacity of forest species to persist will depend on the combination of three non-mutually exclusive processes: migration toward climatically more favorable areas, phenotypic plasticity, and genetic adaptation to new environmental conditions [4]. Given that the rate of population migration is usually lower than the rate of climate shift, the capacity of populations of forest tree species to persist will largely depend on their ability to respond *in situ* through phenotypic plasticity or genetic adaptation [5]. While plasticity is recognized as a key mechanism for responding to rapid environmental changes, medium- and long-term persistence will also depend on adaptation through natural selection in response to new climatic conditions [6, 7].

Adaptive responses of tree populations to environmental changes have been largely studied through common garden experiments [8–10]. By growing populations of different origins under the same environmental conditions, the observed phenotypic differences can be ascribed to genetic effects [11]. In addition, analyzing the associations between population trait values measured under a common environment and the climate of origin of the populations provides evidence of the potential climatic drivers associated with divergent selection along environmental gradients [12–14]. In general, understanding the prevalence of population-level differentiation and the nature of trait-climate and trait-trait associations can help us understand the evolutionary trajectories along past climatic changes. Furthermore, climate patterns of population differentiation can ultimately be used to draw inferences about past evolutionary dynamics [4, 15, 16].

However, adaptive differentiation among climatically contrasting extant populations has built up since their establishment, and does not necessarily mirror temporal adaptive evolution that is of concern under ongoing climate change. Hence, comparing patterns detected in provenance trials with allochronic approaches that allow to analyze the genetic changes over time within the same population may be especially useful for evaluating whether the spatial clines inferred from common gardens with contrasting populations are consistent with adaptive evolution in response to intrapopulation climatic changes [17–19]. However, allochronic genetic studies are extremely rare in trees compared to synchronic studies (i.e. provenance trials) due to time constraints imposed for monitoring temporal changes in long-lived species. Although the comparison suffers from a significant imbalance between the number of synchronic and allochronic studies, it provides key insights into the current debate on the rate of contemporary tree evolution in response to climate change.

In this review, we will revisit trait responses of tree populations detected in provenance trials in light of the underlying temporal dynamics (evolutionary rates) that have shaped population differentiation. In addition, we will compare present population differentiation that has built up during the Quaternary (synchronic approach, *sensu* Hendry and Kinnison [20]) with temporal evolutionary changes over shorter timescales that pave contemporary evolution (allochronic approach). Here, we focus on oaks (*Quercus* sp.) as a study system for several reasons. First, a considerable number of common garden experiments has been conducted across a relatively high number of species from climatically contrasting biomes. This provides a good opportunity to evaluate how adaptation to climatic variability occurs and to identify the main climatic drivers of these processes. Second, oaks show high functional diversity within a single genus, in which labile evolution has been demonstrated for many traits important for adaptation, such as growth form, leaf habit, and traits related to drought and cold tolerance [21– 23]. Third, there are unique allochronic studies that have evaluated recent genetic changes in traits potentially important for adaptation to climatic change, allowing us to assess how these adaptive processes have occurred and whether the patterns are comparable to those found when studying population variation (e.g. [17, 18]).

We address the following specific questions: i) How frequent is population variation in potentially climate-adaptive traits in oak species? ii) Based on common garden studies, how oak populations adapted to climatic gradients? iii) Are long-term patterns of population variation observed in common garden experiments consistent with short-term population changes observed in longitudinal monitoring? iv) Are rates of contemporary evolution comparable to trends of trait divergence?

## 2. Prevalence of population differentiation and genotype-by-environment interaction in oaks

Common garden experiments have been one of the main tools used to study how tree populations genetically differentiate along environmental gradients. We first address the question of how common population variation in traits and plasticity (i.e. G×E interaction) is within the genus by implementing a meta-analysis of studies conducted in common gardens.

For this purpose, we performed a literature search and conducted a meta-analysis of proportions on the resulting studies. The literature search was carried out in the Web of Science (ISI) and Google Scholar looking for population studies conducted under common garden conditions in oak species from all over the world. We used the following keywords and combinations of them: ‘population-level’, ‘population differen*’, ‘population divergence’, ‘population differentiation’, ‘population’, ‘provenance’, ‘local adaptation’, ‘common garden’, ‘trial’, ‘reciprocal transplant’, ‘Quercus’. We applied no date restrictions to the search. We did not consider studies that used populations originating outside of the species’ natural range (e.g. studies on *Quercus rubra* with provenances from Europe). For each study, we recorded the species, the number of traits measured, the number of populations, and the number of experimental sites. We also recorded the number of traits exhibiting significant genetic differences among populations. If a given article evaluated population differentiation in more than one study species, we recorded the information for each species as separate study cases.

Overall, the database comprises 698 observations from 88 studies with several of them including more than one species. Therefore, the total study cases was 96. These studies covered 21 species distributed across three biomes (Figure 1). Specifically, 42.3 % (n = 41) of the study cases were conducted in woodland/shrublands according to Whittaker’s biomes that largely correspond to Mediterranean-type ecosystems, 47 % (n = 45) of the study cases were conducted in temperate seasonal forests, and the remaining 10.4 % (n = 10) in tropical seasonal forests (Figure 1). Geographically, 70.5 % of the study cases (n = 62) focused on species distributed in Europe and/or North Africa whereas only 22.7 % (n = 20) were conducted in species from America, where the genus has its highest diversity, and 5.7 % (n = 5) in Asia.

**Fig 1.**
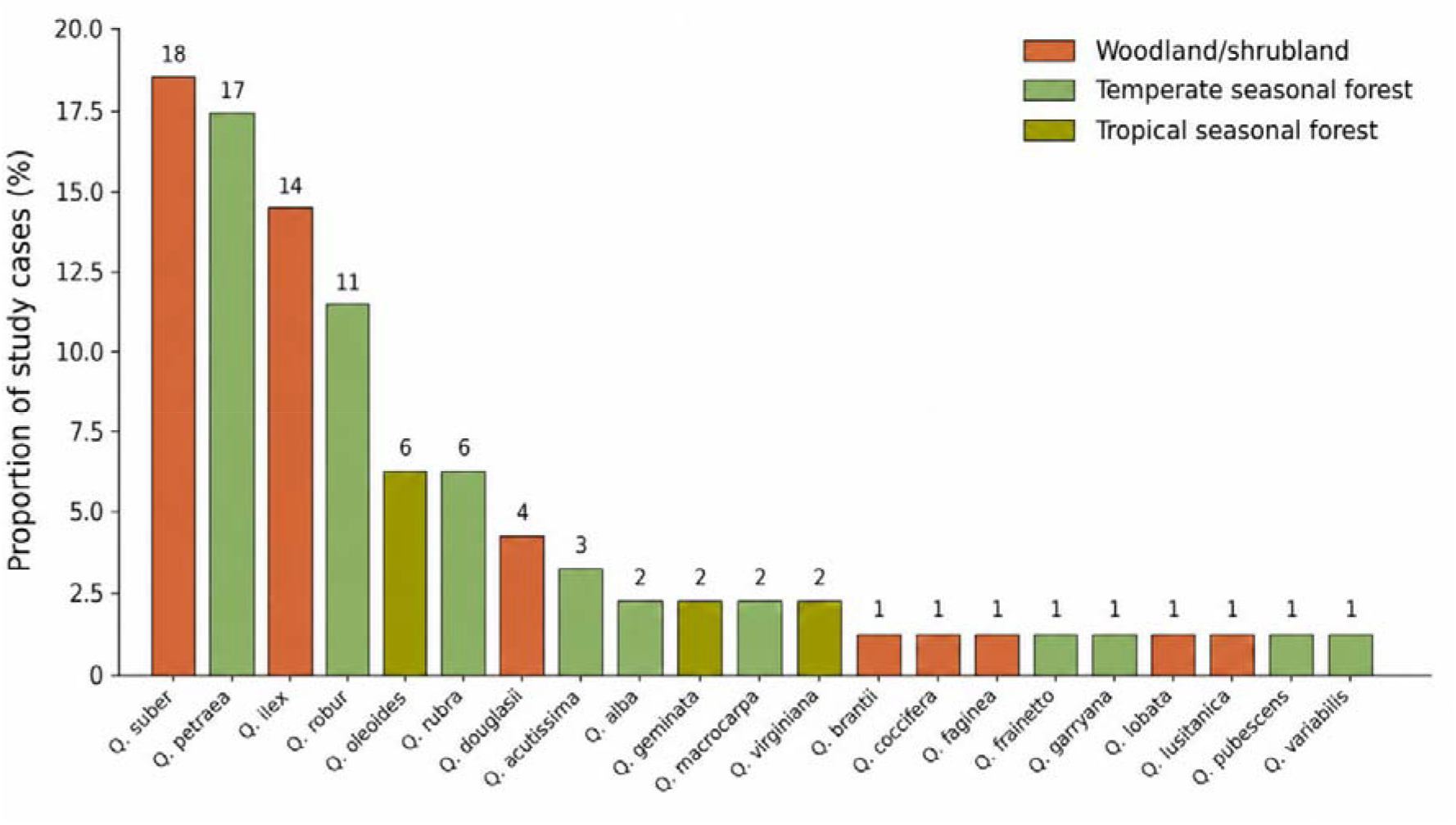
Proportion of study cases per species, expressed as a percentage. The total number of studies was 88, but several studies included more than one species, so the sum used to estimate the percentage of study cases was 96. Numbers above bars indicate the number of study cases for each species. Colors indicate the main biome in which each species is distributed, as shown in the legend.

We estimated the overall frequency (i.e. proportion) of traits reporting significant (*P* < 0.05) population genetic differentiation by conducting a meta-analysis of proportions [24] using the *metafor* package [25] in R 4.4. Briefly, to do that, we related the number of traits showing significant population differentiation to the total number of traits evaluated for each study case. Proportions were transformed using the arcsine square-root transformation, and their corresponding sampling variances were calculated using the *escalc* function. We then fitted a random-effects meta-analytic model using rma.mv, with study identity and species included as random effects to account for the non-independence of observations from the same study or species.

The results of our meta-analysis revealed that, on average, population genetic differentiation in the genus is common and widespread with 61.54 % (95% CI 42.58–78.84, P < 0.001) of the traits showing significant population differentiation. An Egger meta-regression detected significant funnel-plot asymmetry (Z = 4.58, P < 0.001). Since the sampling variance of each proportion depended on the number of traits evaluated in each study case, this result likely reflect a relationship between the estimated proportion and the number of traits assess. The results of the meta-analysis indicated significant population variation across studies in oaks although at a lower frequency than reported in previous reviews of forest tree species conducted in other phylogenetic groups or geographical regions. For example, Aspalter et al. [26] found a prevalence of 73 % of population differentiation in European forest tree species, Leites & Benito-Garzón [14] found evidence of local adaptation in 79 % of the species studied, with evidence being more common in conifers (87.5%) than in broadleaf species (67%). Ramírez-Valiente et al. [13] found that 82.9 % of the studies in Mediterranean species (where 80% of the studies were conducted in *Pinus* sp.) exhibited genetic differences among populations. Alberto et al. [12] in a review on provenance trials mainly conducted in temperate regions found that 89.8 % of the studies exhibited significant differences among populations. These results indicate that most studies on oaks show significant population-level variation but to a lesser extent than other taxonomic groups.

The meta-analysis also revealed high heterogeneity across studies (heterogeneity test Q = 818.4, P < 0.001), indicating that population genetic differentiation is highly variable across studies. We tested to what extent this heterogeneity resulted from differences in biomes, phylogenetic sections, leaf habit, life stage and type of functional trait (e.g. growth, gas exchange, phenology, etc.) by including them as moderators in the meta-analysis. We found that biome (QM(df = 2) = 5.70, P = 0.058), phylogenetic section (QM(df = 4) = 3.64, P = 0.457), leaf habit (QM(df = 1) = 3.65, P = 0.056) and life stage (QM(df = 3) = 6.68, P = 0.087) were not significant. In contrast, functional trait category (QM(df = 27) = 160.48, P < .0001) was highly significant. These results indicate that differences among functional traits (categories) were important factors explaining differences in population genetic variation across studies. In other words, population genetic differences change across studies as a result of measuring different functional traits. In general, traits related to growth, phenology, leaf morphology and allometry exhibited high population differentiation (data not shown), similar to the results obtained by Aspalter et al. [26]. These results do not necessarily indicate that some traits are more important than others for adaptation, but rather that these are the traits in which greater population differences are detected across studies, which may be due to several reasons. Research focus bias is likely one of them, as some traits are more frequently investigated generally because they are easier to measure, or receive more attention for economic reasons. Some functional traits that are potentially important for adaptation to stresses, such as osmotic adjustment, resistance to xylem cavitation, or tolerance to freezing temperatures, have been studied in too few studies to be tested in this analysis. The high phenotypic plasticity of some traits such as those related to gas exchange or photochemistry may allow many species to adjust them in response to environmental variation, potentially limiting divergent natural selection (e.g. [27, 28]).

Finally, 27.0 % (13.3-43.4, P < 0.001) of traits within oak studies also showed significant population-by-environment interaction, indicating differences in phenotypic plasticity among populations. This proportion is also lower than that reported in previous reviews on European trees [26], Mediterranean species [13] and plants in general [29].

### 3. Adaptation patterns of oak populations across biomes

In this section, we reviewed studies on common gardens to better understand how different species have adapted to climatic gradients across different biomes and biogeographic regions worldwide.

### 3.1. Adaptation patterns in seasonally temperate forests

In species from seasonally temperate forests, resource-use strategies appear to be associated with temperature and precipitation gradients. In general, the literature shows a tendency for populations from warmer sites to have leaves with lower SLA, higher thickness, longer growing seasons through earlier spring phenology and/or later leaf senescence, and higher growth potential under favorable conditions. In addition, these traits are closely associated in several studies in *Q. petraea, Q. robur, Q. alba, Q. macrocarpa, Q. ellipsoidalis*, and *Q. variabilis* suggesting to be part of a functional syndrome or strategy [30–34].

Specifically, regarding leaf morphology, Torres-Ruiz et al. [30], in a study with ten populations of *Q. petraea*, found that populations from warmer and more mesic areas exhibited lower SLA (Fig. 2). In *Q. rubra*, Gómez-Quijano et al. [35], in a study with eight populations, also found that populations from sites with higher mean temperatures and higher summer precipitation had lower SLA. Etterson et al. [31] found very similar patterns for both *Q. rubra* and *Q. macrocarpa*, for which populations from lower latitudes with warmer climates also showed lower SLA, a more extended growing season, and greater height growth. These patterns of low SLA in populations from warm and humid sites, associated with higher growth, contrast with the Leaf Economics Spectrum hypothesis, in which higher SLA is expected to be associated with higher mass-based photosynthetic rates and higher relative growth rates [36, 37]. These patterns are also opposite to those found in Mediterranean regions (see the following section), where low SLA is generally associated with drier climates and has been interpreted as an adaptation conferring desiccation tolerance and longer leaf life spans in poorer environments [38–40]. Some authors suggest that lower SLA may be related to a longer leaf life span, associated with a strategy of longer growing-season duration and, consequently, higher growth potential [41].

**Fig 2.**
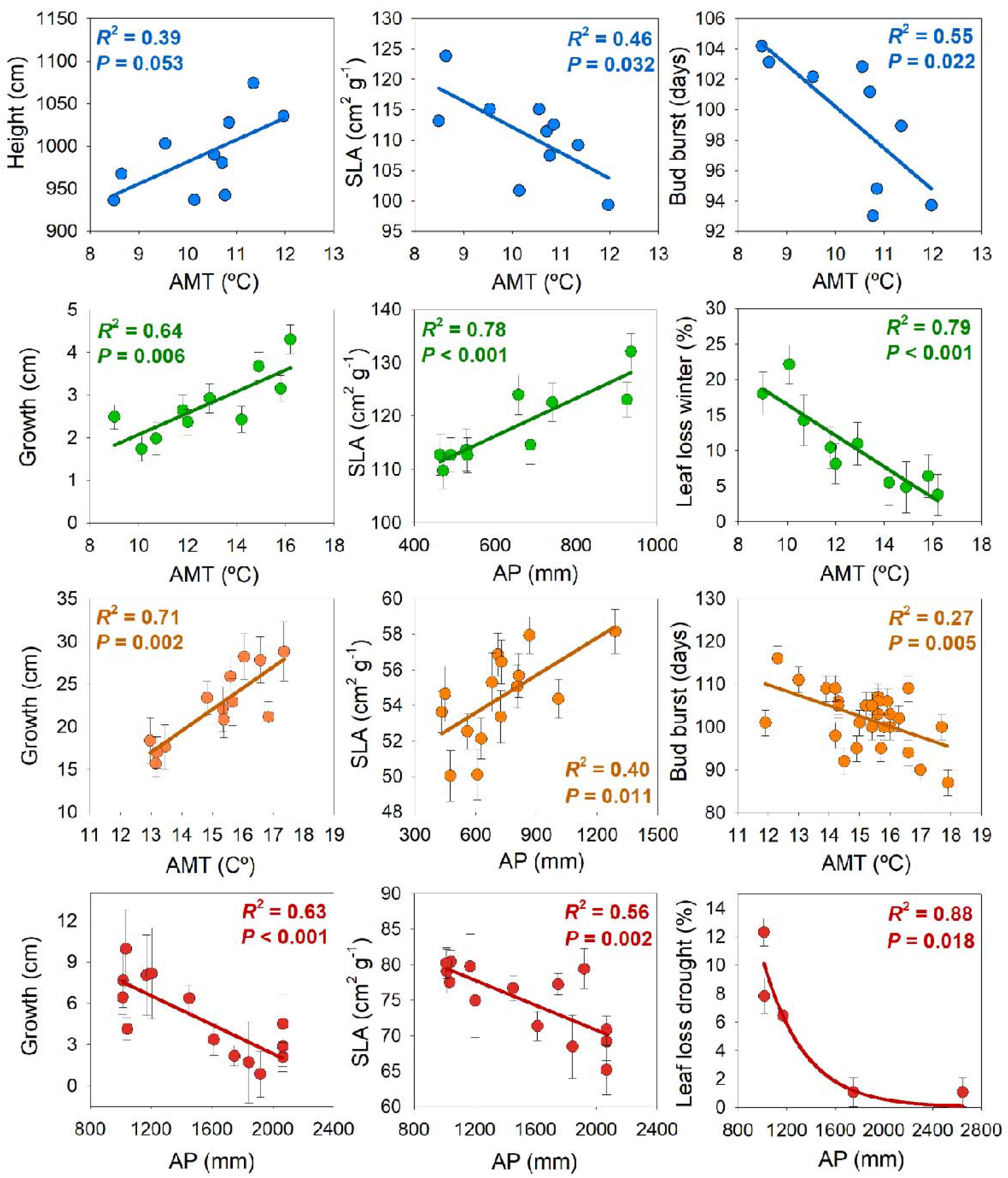
Representation of the associations between source-population climate, defined by mean annual temperature (AMT) and annual precipitation (AP), and growth, SLA, and phenological traits measured in common gardens. The figure shows patterns for four oak species, *Q. petraea* (blue), *Q. faginea* (green), *Q. suber* (orange), and *Q. oleoides* (red), representing the three main biomes in which oaks are distributed: temperate seasonal forest, woodland/shrubland, and tropical seasonal forest. Within woodland/shrubland, two species with contrasting leaf habits, deciduous (*Q. faginea*) and evergreen (*Q. suber*), are represented. Data were extracted from Torres-Ruiz et al. [30] for *Q. petraea*, Solé-Medina et al. [40] for *Q. faginea*, Ramírez-Valiente et al. [38] and Sampaio et al. [86] for *Q. suber* and Ramírez-Valiente & Cavender-Bares [28] and Ramírez-Valiente et al. [93] for *Q. oleoides*. Growth is represented by height growth in Q. petraea and annual growth for the rest of the species. For simplicity, a single species and values from a single common garden are shown as an example of the most general trends observed in these biomes.

Wang et al. [34], in a study on *Q. variabilis*, an Asian species distributed across temperate regions, found that populations from warmer and more mesic sites had higher leaf thickness, similar to the results from European and American species. Wang et al. [34] also analyzed a set of traits related to leaf anatomy. Specifically, they found that populations from warm and humid sites had a thinner palisade mesophyll, a thicker spongy mesophyll, smaller and denser stomata, and wider vessels. A more developed spongy mesophyll may favor internal CO_2_ diffusion and increase leaf hydraulic capacitance, whereas a higher density of small stomata may support higher gas exchange capacity and faster stomatal responses [42–44]. Likewise, wider vessels would increase hydraulic efficiency, and thicker vessel walls could provide mechanical reinforcement against conduit collapse under negative xylem pressure [45, 46]. Therefore, the pattern of thicker leaves with lower SLA in warmer and wetter areas could be related to greater hydraulic and gas exchange capacity.

Population differences in phenology also appear to be associated with clines although the direction of these clines differs between Eurasian and North American species. In Eurasian species, populations from warmer and more mesic sites generally show earlier spring budburst and/or later fall senescence (Fig. 2). Such clines have been repeatedly observed in common gardens in Eurasian species such as *Q. mongolica* [47], *Q. petraea* [30, 48–54], and *Q. robur* [55]. In North American species, by contrast, the spring phenology is often delayed in warmer or more southern provenances, contrary to the pattern observed in Eurasian species [31–33, 56–58]. Interestingly, although colder and northern populations tend to flush earlier, warmer and lower-latitude provenances also retain their leaves for longer because of later senescence. Thus, extended growing seasons in temperate oaks can arise through different phenological routes: earlier spring flushing and delayed senescence, and sometimes a later-shifted growing season, particularly in several North American species.

Opposite genetic clines between North American and Eurasian species in spring phenology likely reflect different selective pressures acting on the timing of bud burst on both continents. The climate of eastern North America is characterized by a higher frequency of extreme cold events after warm spells in spring, exposing trees to more frequent late frost events [19]. As a result, selection may favor different combinations of phenological traits: late flushing as a frost-avoidance strategy in some populations, but also early flushing coupled with cold tolerance in northern populations adapted to short growing seasons. This may explain why North American provenances from colder regions can flush earlier while still avoiding or tolerating frost damage more effectively than southern provenances when planted near the northern range limit.

Together, the studies in temperate oaks show a close relationship between SLA, leaf life span, phenology and growth (Fig. 2), suggesting that the higher growth potential of populations from mesic and warm sites is associated with an extended growing season, longer-lived leaves, and higher potential for gas exchange [30–31, 54]. However, the positive relationship between extended leaf phenology and growth is context dependent. In some species, some populations from warmer sites show lower growth as observed in *Q. petraea* [59], *Q. robur* [60], and *Q. rubra* [33] in some common gardens with significant population-by-environment interaction. Vitasse et al. [51] suggest that a longer growing season may increase exposure to severe spring or autumn frosts, which can reduce annual growth and survival in cold areas where frost risk is higher. In addition, temperature-driven genetic clines may be obscured in common gardens with limited geographic sampling as observed in some species where studies with a higher number of populations have detected adaptive patterns across temperature gradients [61–65].

Drought intensity also appears to be a significant climatic driver for temperate species, although not as generalized as temperature. This has been evidenced in several species such as *Q. robur* and *Q. petraea*, where populations from drier sites show greater growth capacity and survival in drier common gardens [59, 66, 67]. Kubiske and Abrams [68] also found that, in *Q. rubra*, individuals originating from a xeric site had thicker leaves, lower SLA, smaller leaves, and a higher capacity to maintain positive net photosynthesis at lower water potentials, as well as lower non-stomatal inhibition of photosynthesis under drought, than individuals from a mesic site. In *Q. petraea*, Rabarijaona et al. [69] found that populations originating from drier areas had a higher capacity to increase WUEi under drought and exhibited a smaller reduction in radial growth, suggesting that physiological plasticity may contribute to growth stability under stress. Nosenko et al. [65], in a study comparing German populations of *Q. robur* in a common garden with contrasting watering treatments, found that high constitutive iWUE increased in populations originating from regions with lower water availability. In addition, the authors found that climatic clines in δ^13^C appeared mainly under extreme drought, rather than under control conditions, suggesting that adaptive differentiation may become visible only when plants are exposed to sufficiently strong water stress.

### 3.2. Adaptation in woodland/shrublands (Mediterranean-type ecosystems)

Mediterranean regions are characterized by summer drought, a period during which precipitation is lower than potential evapotranspiration. The length and intensity of this dry period vary considerably across regions. In these ecosystems, precipitation, both annual and summer, appears to be a key driver of population genetic divergence in traits related to drought responses. One of the trait categories showing the strongest differentiation among populations across studies is leaf morphology (Fig. 2). In particular, in both deciduous and evergreen species, populations from drier areas generally exhibit more sclerophyllous leaves with lower specific leaf area (SLA) and higher leaf thickness in species such as *Q. suber, Q. ilex, Q. faginea* and *Q. lusitanica* [38–40, 70–77].

Several lines of evidence support the adaptive value of lower SLA under dry conditions. Leaves with lower SLA exhibit thicker external epidermal walls and higher cellulose concentrations, conferring greater resistance to cell collapse, loss of mechanical integrity during dehydration, and higher resistance to elevated temperatures and radiation [41]. In addition, studies using phenotypic selection analyses in Mediterranean oaks have reported that individuals with lower SLA exhibit higher fitness under drier conditions, suggesting an adaptive advantage (e.g. [38, 39, 78]). Furthermore, SLA is commonly associated with other traits related to drought tolerance, including physiological and allometric traits. For example, in *Q. suber*, Ghouil et al. [76] in a study including populations from the southern edge of the distribution range, showed that populations from more xeric areas were characterized by lower SLA but also with smaller leaves and higher elastic adjustment. Ramírez-Valiente et al. [38] also showed in this species that populations with lower SLA exhibited higher water-use efficiency (WUE), measured by 13-Carbon isotope composition (δ^13^C), particularly during drier years. In *Q. ilex*, Peguero-Pina et al. [77, 79] found that the driest populations, grouped within subspecies ballota, exhibited a more negative turgor loss point, indicating a greater capacity to maintain turgor at low water potentials, a higher modulus of elasticity, stiffer cell walls and higher resistance to tissue deformation during dehydration, and higher leaf specific hydraulic conductivity. Together these results suggest higher tolerance to drought in these populations. In *Q. faginea*, under drought, populations with lower SLA also exhibited greater investment in root biomass and higher photochemical efficiency, suggesting that these populations were able to maintain a better physiological status [40]. Similarly, in *Q. douglasii*, Poudel et al. [80] found that seedlings from drier sites maintained higher maximum photochemical efficiency, a higher proportion of green leaves and higher stomatal conductance under drought. In general, several studies have reported that populations originating from drier sites have evolved towards a higher investment in roots [40, 75, 81, 82], which confers overall greater survival under stronger water stress [13, 71, 73].

However, not all studies show such clear population differences in traits related to drought responses, or these differences are not aligned with climatic gradients in precipitation. For example, Gimeno et al. [27] found significant phenotypic plasticity in physiological traits such as photosynthetic rates or water use efficiency in response to water availability in six *Q. ilex* populations, but did not detect population differences, suggesting that physiological plasticity may override the population differences. In the same species, Juan-Ovejero et al. [83] also found no ecotypic differentiation among four populations in leaf 18-Oxygen isotope discrimination, nutrient traits, leader shoot length or aboveground biomass growth. In contrast, intrapopulation variation was considerably higher, suggesting that much of the variability of the species is contained within populations. In *Q. douglasii*, Skelton et al. [84] reported low population differentiation in vulnerability to embolism, similar to Peguero-Pina et al. [79] in *Q. ilex*, despite having found significant differences in many other traits (see comments above), suggesting that some hydraulic traits may be more canalized or may show adaptation at the tissue level rather than among populations.

On the other hand, many studies show population differences in growth, but unlike what occurs for most drought-related traits, population variation appears to be more strongly associated with temperature gradients (Fig. 2), with populations from warmer sites and lower latitudes showing higher growth under favorable conditions [38, 40, 75, 82, 85–87]. Phenological traits and those associated with cold responses also show population variation associated with temperature clines. For example, Sampaio et al. [88] found that populations regions of *Q. suber* originating from areas with higher annual mean temperature exhibited earlier bud burst, suggesting that part of the variation in growth may be mediated by differences in the length of the growing season, as occurs in temperate species. Meanwhile, Aranda et al. [89], also in *Q. suber*, showed population differences in responses to low temperatures, with populations from colder sites exhibiting lower winter photoinhibition than populations from warmer sites. In *Q. faginea*, a marcescent species, Martín-Clar et al. (unpublished) show that populations from sites with warmer winters, which also grow significantly more, retain a higher proportion of leaves during winter (Fig. 2). Together, these patterns suggest that in regions with mild winters, selection has favored a strategy of exploiting thermally favorable windows, particularly during winter and spring, by maintaining carbon fixation for longer periods, initiating growth earlier or retaining leaves for longer. In contrast, in continental regions or colder environments, populations have evolved towards increased tolerance to freezing temperatures and delayed phenology that avoids damage from late frosts, reducing resource allocation to growth.

### 3.3. Adaptation in tropical biomes

In the few oak species from tropical seasonal forests for which common garden data are available, patterns of population differentiation appear to be mainly associated with precipitation gradients rather than temperature [21, 90, 91]. This is particularly evident in the neotropical oak *Quercus oleoides*, a species widely distributed across Central America along a marked precipitation gradient, with a dry season lasting between 2 and 6 months. Specifically, one of the most striking patterns is that population variation in leaf morphology contrasts with that observed in Mediterranean species (Fig. 2). As discussed in the previous section, the adaptation of populations of Mediterranean species to drier sites has generally involved a conservative strategy with more sclerophyllous leaves with low SLA and traits associated with drought tolerance, such as more negative water potential at the turgor loss point. In *Q. oleoides*, however, populations from more xeric origins with long and intense dry seasons, tend to produce larger, thinner leaves with higher SLA, whereas populations from more mesic origins develop smaller, thicker, more sclerophyllous leaves with lower SLA (Fig. 2). This pattern has been observed in multiple common gardens with different populations from contrasting geographic ranges [21, 28, 92–94].

This pattern, which may seem counterintuitive, can be understood when other traits related to drought responses and growth potential are analyzed. In *Q. oleoides*, populations from more xeric sites show leaf abscission in response to drought, suggesting a partial drought avoidance strategy based on the seasonal reduction of the transpiring surface area (Fig. 2) [21, 28]. By contrast, populations from more mesic origins, where the dry season is shorter or less severe, have leaves with lower SLA, as well as smaller area and a significantly higher capacity for osmotic adjustment than xeric populations, such that they are able to decrease the turgor loss point to a higher extent. This indicates a higher capacity to maintain leaf turgor, and therefore gas exchange, at more negative water potentials [21, 28]. Likewise, more mesic populations also show greater activation of photoprotection associated with the xanthophyll cycle under drought, indicating that leaves that remain physiologically active during water stress increase the capacity to dissipate excess light energy [90]. In addition, Ramírez-Valiente et al. [92], in a field common garden study, showed that populations from environments with longer and more severe dry seasons exhibited higher growth rates during the wet season, but not during the dry season, supporting the idea of an acquisitive strategy during favorable water conditions coupled with an avoidance strategy under drought.

Together, these results suggest that, in tropical seasonal environments, under more mesic conditions, natural selection favors the evolution of more sclerophyllous leaves and higher osmotic adjustment, possibly as a response to tolerate a shorter dry season while maintaining higher carbon assimilation under this period. If the dry season becomes more intense and prolonged, a strategy based on reducing the transpiring surface area through less costly leaves with greater carbon-fixation potential during the wet season may be beneficial. This pattern contrasts with that observed in Mediterranean environments, probably because in the latter there is a conjunction of two important selective factors, summer drought and low winter temperatures. Indeed, studies of xeric populations at the southern range margins of Mediterranean species, in subtropical climates lacking a period of winter stress, show that populations tend to have less sclerophyllous leaves [94, 95]. This observation could be associated with an acquisitive and drought avoidance strategy similar to that observed in seasonally dry tropical oaks. Thus, it would be important to increase the number of studies at the southern distribution limits of Mediterranean species, and in a larger number of species from tropical regions, including summer deciduous species, to better understand how species adapt to drought in the absence of the constraints imposed by a period of winter cold.

## 4. Population evolution

In contrast to synchronic approaches, which dissect population genetic differences among present-day spatially separated populations, allochronic approaches track temporal genetic (evolutionary) changes within the same population. While an extensive body of results is available from synchronic studies, allochronic case studies remain scarce and are largely limited to very few species (Fig. 3).

**Fig 3.**
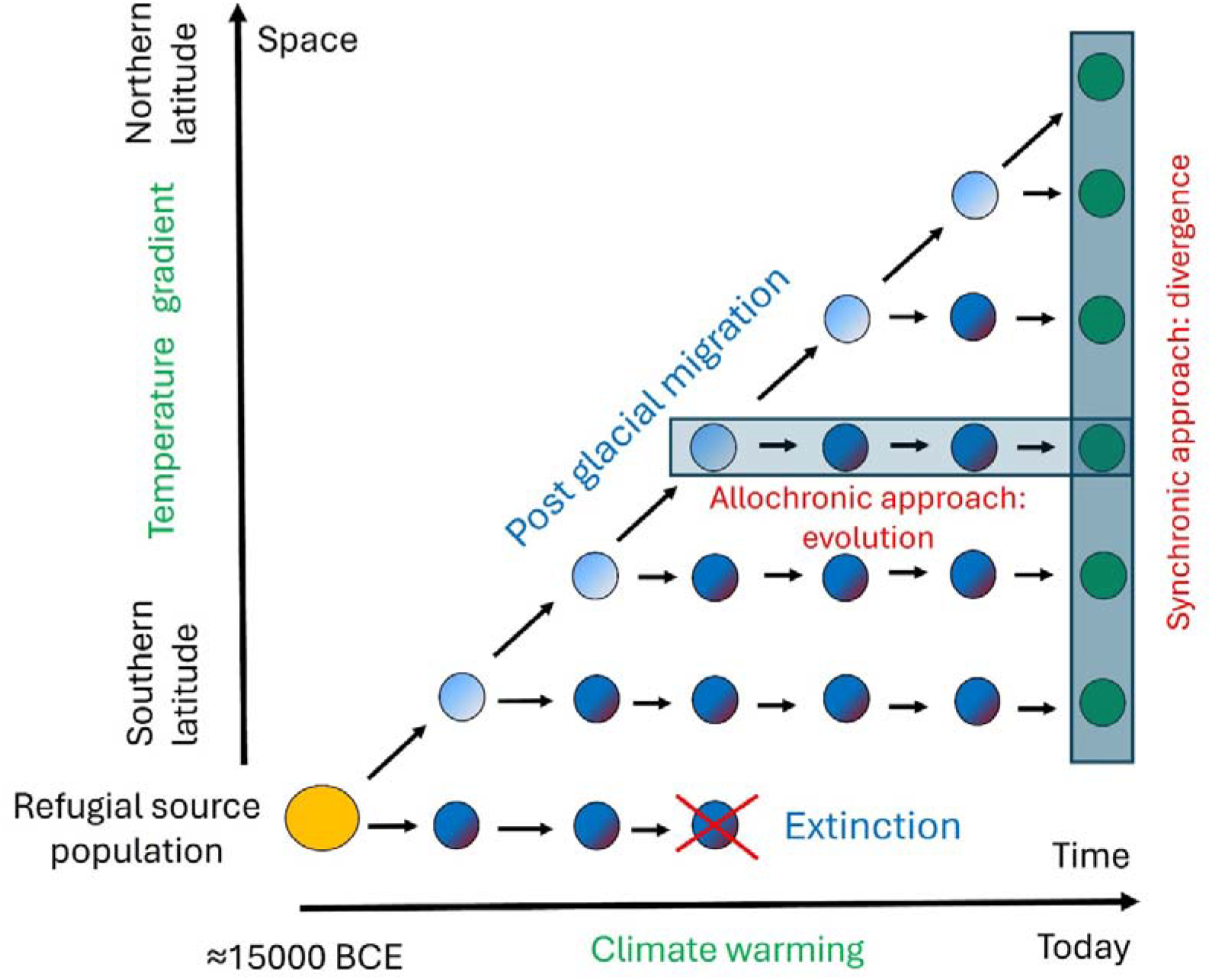
Synchronic and allochronic approaches used to study microevolutionary changes in trees. This figure sketches the evolutionary trajectory of European temperate oak populations (blue circles) since the Last Glacial Maximum (LGM) in Europe and illustrates the allochronic and synchronic approaches implemented for assessing evolutionary changes. As the temperature steadily increased during the Holocene, oaks migrated northwards from the refugial areas located in Southern Europe and founded new populations along a latitudinal gradient (black arrows). Extinctions were limited in the southern areas. Both the spatial and temporal environmental gradients were dominated by temperature variation, from south to north and from the LGM to the present. Extant populations (vertical underlined column) sampled in common garden experiments (synchronic approach) exhibit genetic differentiation that have accumulated since their divergence from the refugial populations along the spatial and temporal gradients. In contrast, allochronic approaches focus on the temporal genetic changes in individual populations across successive generations that have paved recent or ongoing temperature changes (horizontal underlined row as an example).

Tracking temporal genetic changes within the same population across generations (longitudinal monitoring) remains a challenging task because of biological constraints related to the long lifespan of oak trees. Since longitudinal monitoring can hardly be implemented over more than two generations, historical and retrospective monitoring offer attractive alternatives for allochronic approaches. Historical monitoring aims to compare trait values in present-day populations with those of ancestral populations, inferred from evolutionary models or coancestry reconstructions [96, 97]. Retrospective monitoring consists of assessing traits in age-structured cohorts of living trees. Most oak stands are composed of overlapping generations, including trees belonging to different age-structured cohorts. Assuming that all cohorts derive from the same recent ancestral population, differences among cohorts may reflect temporal genetic variation [17, 18]. Indeed, the demographic dynamics of oak stands indicate that the strongest evolutionary events during stand development occur at very early stages, when most seedlings are eliminated through natural selection, competition, and stochastic mortality. Thus, a present-day cohort that is approximately 100 years old mainly reflects evolutionary changes that occurred about 100 to 90 years ago. This constitutes the rationale for using retrospective monitoring to assess temporal genetic changes in oaks.

Evidently, generational links are lost in historical and retrospective monitoring compared with longitudinal monitoring. By definition, longitudinal, retrospective, and historical monitoring encompass different temporal scales, ranging from a few decades for longitudinal monitoring to centuries for retrospective monitoring and millennia for historical monitoring. Oak-based examples illustrating all three approaches provide some of the first insights into evolutionary rates estimated from allochronic studies.

### 4.1 Evolution during the Quaternary (historical monitoring)

The method developed by Ovaskainen et al. [96] and Karhunen et al. [97] enables the retrospective reconstruction of functional trait evolution using phenotypic data from common gardens and genomic information from the populations. Briefly, this method is based on the construction of a coancestry matrix among populations using genomic data, under a model in which current populations derive from a common ancestral gene pool and have partially diverged through genetic drift and gene flow. Second, this matrix is incorporated into an animal model [98] that uses phenotypic information of a genetic relatedness structure obtained from common gardens to estimate the additive genetic values of the populations. In addition, the model estimates the additive genetic mean of a hypothetical ancestral population, which acts as an evolutionary reference against which current populations can be compared.

To include the temporal dimension of this evolutionary change, coalescent-based demographic models can be used [99, 100]. Thus, the integration of both methodologies allows the estimation of evolutionary rates of traits per generation. Although these estimates should be interpreted with caution, especially when gene flow or uncertainty in divergence times exists, they provide a way to estimate temporal changes in traits throughout the evolutionary history of species. This framework is particularly useful in forest trees, where direct allochronic studies are scarce because of their long generation times, but where common gardens and available genomic resources enable to infer the historical evolutionary trajectories of current populations. Using the data from Ramírez-Valiente et al. [72], we applied these methods to estimate evolutionary rates of functional traits in the Mediterranean *Q. faginea*. Briefly, in this study, eight natural populations of *Q. faginea* were genotyped using 11,463 SNPs and the same populations were established in a common garden, where functional characterization was carried out for eleven traits. We estimated divergence times between each pair of populations and their closest common ancestors using coalescent models implemented in FastSimCoal2 [101]. We also estimated the additive genetic values of traits for the populations and their closest common ancestors using the driftsel R package [102]. For each pair of populations, alternative divergence models were compared: divergence in strict isolation, divergence with symmetric migration, and divergence with asymmetric migration.

The most recent divergence times estimated by FastSimCoal2 were 1139 generations for the VAL-SAL populations, 1257 generations for the VAL-SOT populations, and 1730 generations for the BOS-PEN populations (Table S1, Fig. 4). In all three cases, the best-supported model was isolation with asymmetric migration, indicating that the recent demographic history of these populations cannot be interpreted as a simple split in isolation, but rather as divergence accompanied by gene flow among populations (Table S1, Table S2). Using these divergence times, we calculated evolutionary rates as the percentage change relative to the ancestral value, divided by divergence time. Evolutionary rates were highest for the branches leading to VAL and PEN populations (Fig. 4). For SLA, the trait showing the highest evolutionary rates (data not shown), we found a rate of change of +0.004275% generation^−1^ for VAL from its common ancestor with SAL, +0.005884% generation^−1^ for VAL from its common ancestor with SOT, and +0.003945% generation^−1^ for PEN from its common ancestor with BOS. By contrast, SAL, SOT, and BOS showed negative rates, indicating relative decreases in SLA with respect to their respective common ancestors (Fig. 4).

**Fig 4.**
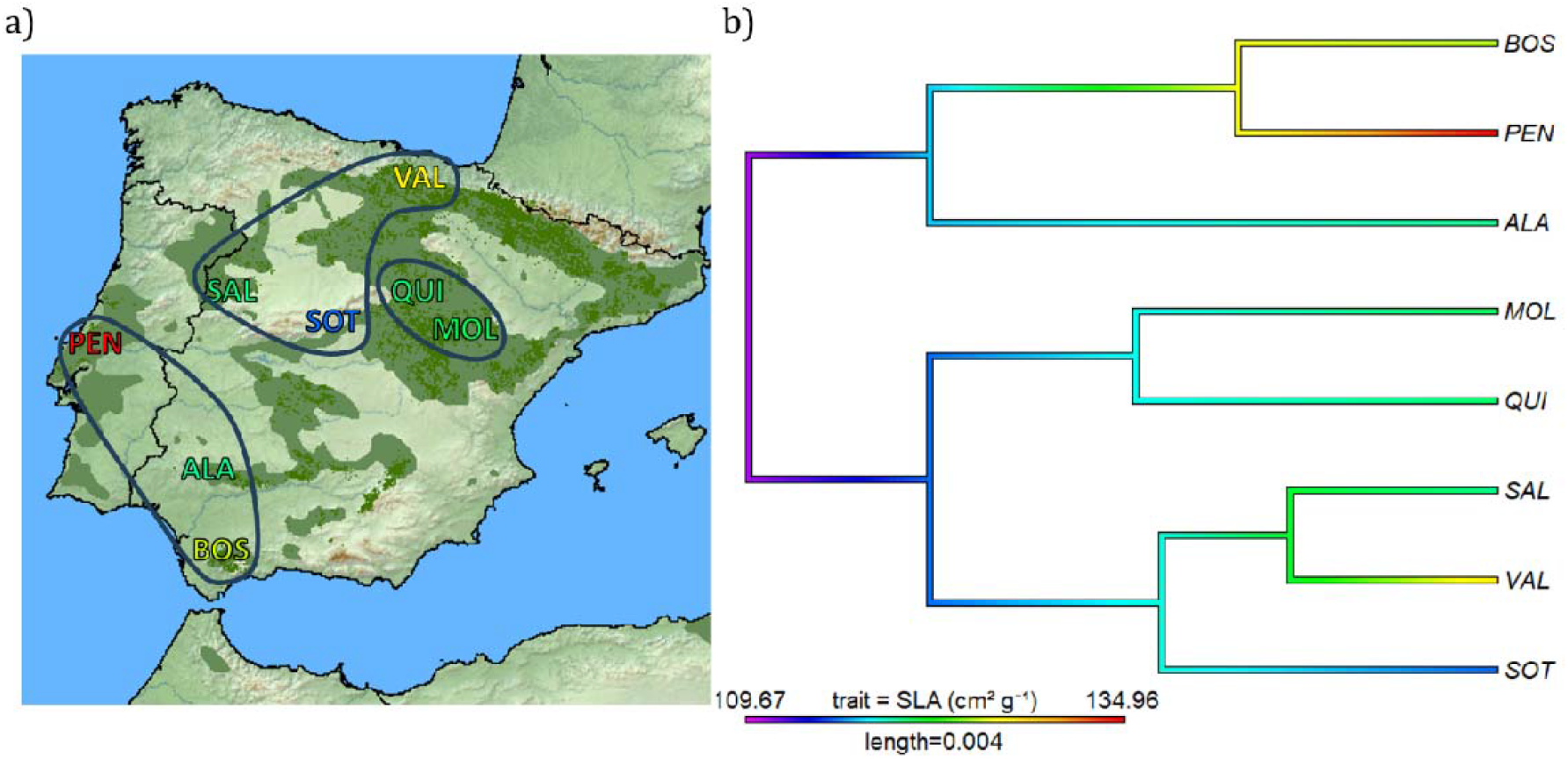
Estimation of evolutionary rates in *Q. faginea*. a) Populations of *Q. faginea* for which genomic analysis using dd-RADseq and phenotyping under common garden conditions were conducted in Ramírez-Valiente et al. [72]. The green area shows the distribution of *Q. faginea* in the Iberian Peninsula. Population colors correspond to SLA values shown in b. Blue lines group the genetically closest populations according to Ramírez-Valiente et al. [72]. b) Dendrogram of genetic distances among the six populations of Q. faginea, constructed using the unweighted pair group method. Colors indicate the evolution of the additive genetic value of the target trait, SLA, from the hypothetical ancestral population to the current populations, following the method of Ovaskainen et al. [96].

The evolutionary reconstruction suggests that trait evolution has not followed a linear and sustained trajectory in a single direction (e.g., SLA, Fig. 4). In some cases, ancestral nodes and current populations show changes of opposite sign along nearby branches. For example, in the case of SLA in the northwestern group of populations, the pattern suggests that some lineages have evolved toward higher SLA values, whereas others have evolved toward lower values. This pattern is consistent with fluctuating evolution or nonlinear evolutionary trajectories, potentially generated by temporal or spatial changes in selective regimes. Therefore, evolutionary rates estimated over long temporal scales are expected to be more conservative than the evolutionary changes resulting from rapid climatic shifts.

### 4.2. Evolution since the Little Ice Age (retrospective monitoring)

Retrospective monitoring has been conducted in three sessile oak (*Quercus petraea*) forests. Within each forest, four age-structured cohorts established at different times between 1680 and the present day (1680, 1850, 1960, and 2008) were compared for various functional traits [17, 18]. The same cohort sampling design was replicated in three forests located in western France (Tronçais, Bercé, and Réno-Valdieu). The three selected forests were managed under even-aged silvicultural regimes, which facilitated the sampling of entire stands (cohorts) composed of trees of approximately the same age and originating from natural regeneration. In the three oldest cohorts (1680, 1850, and 1960), acorns were collected from thirty trees per cohort and grown in a common garden experiment. Seedlings originating from the three cohorts were then compared by assessing functional traits under common garden conditions. The temporal scale considered in this experiment spans the last three centuries and encompasses the transition from a cold climatic period (1680-1850) to a warmer one (1850-1960). The former coincides with the end of the Little Ice Age, which extended from the mid-fifteenth century to the mid-nineteenth century [103], whereas the latter corresponds to the early warming phase of the Anthropocene, beginning during the second half of the nineteenth century. Climatic variations during these periods are well documented through instrumental and historical records [104, 105].

Results from Saleh et al. [17] and Caignard et al. [18] showed that growth traits (height and diameter), phenological traits (timing of bud burst and marcescence), and leaf morphological traits (SLA, specific leaf area) exhibited temporal genetic changes across cohorts closely associated with climatic fluctuations. These changes, which were consistent across the three forests, were characterized by reduced growth during cold periods and enhanced growth during warm periods, earlier bud burst during cold periods, more pronounced marcescence during warm periods, and lower specific leaf area during warm periods.

Importantly, temporal changes fluctuated in parallel with climatic transitions between cold and warm conditions (Fig. 5). This fluctuating pattern is particularly evident during the transition from the Little Ice Age to the onset of the Anthropocene. For example, while growth decreased in all three forests during the cold phase of the Little Ice Age, it increased during the subsequent warming period in the same forests (Fig. 5). Similar fluctuating patterns were also observed for bud burst, with earlier flushing during the cold period in all three forests and later flushing during the warm period in two forests [18]. In summary, short-term fluctuations in temporal genetic variation of growth traits provide strong evidence that evolutionary processes can occur on surprisingly short timescales, even among long-lived species. Moreover, the observed fluctuating evolutionary trends call for a reassessment of long-term trend estimates. Long-term patterns assessed over extended time-periods can obscure short-term underlying changes driven by fluctuating environmental changes. When successive responses shift in opposite directions over short time scales, the overall trend estimated across the entire study period will underestimate the magnitude of the instantaneous changes.

**Fig 5.**
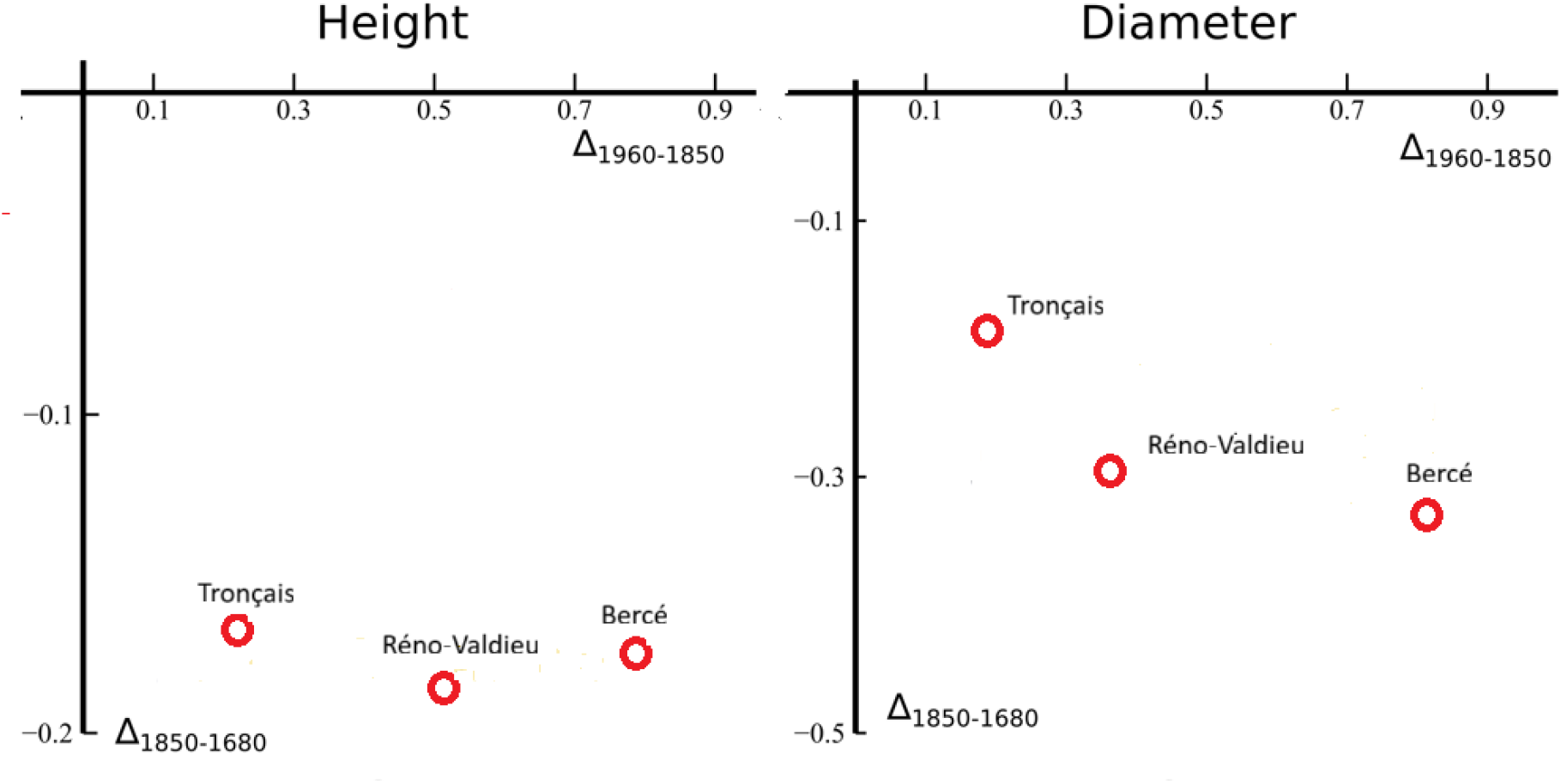
Fluctuating temporal genetic changes of height and diameter in *Q. petraea* between the cold period at the end of the Little Ice Age (1680-1850) and the warming period at the onset of the Anthropocene (1850-1960), assessed in three forests (according to Caignard et al. [18]). The Δ values correspond to the differences between two successive cohorts expressed in standardized values. For example, Δ_1960-1850_ of height stands for the difference in height between 1960 and cohorts 1850 as assessed in offspring from the cohorts raised in a common garden (see text).

### 4.3 Evolution during recent decades (longitudinal monitoring)

Truffaut et al. [106] and Alexandre et al. [107] monitored genetic changes in phenotypic traits across two successive generations in a sessile oak (*Quercus petraea*) stand located in the Petite Charnie forest in western France. The stand was approximately 100 years old (generation G1) when it underwent seed cutting in 1989, followed by secondary cuttings and a final clear-cutting in 2001. Systematic sampling of the subsequent regeneration was then carried out between 2014 and 2017 (generation G2). Genetic relatedness among G1 trees, among G2 seedlings, and between G1 parent trees and their G2 offspring was reconstructed using genetic and genomic fingerprints. Access to these kinship relationships within and between generations made it possible to estimate the genetic values (breeding values) of functional traits in both generations, thereby providing insight into temporal genetic changes. Longitudinal monitoring was conducted on the same traits as for retrospective monitoring. The study period corresponded to a phase of steadily increasing temperatures between 1989 and 2014.

Significant genetic changes between the two generations were observed for growth traits and leaf traits (MLA, mean leaf area, and SLA, specific leaf area) and carbon content of leaves. No changes were detected for phenological traits and other physiological traits. Those genetic changes between to succesive generations are in range of a few percent (Table 1). Comparing the results of retrospective monitoring with longitudinal monitoring, convergent trends can be observed towards increasing growth and decreasing specific leaf area during the warm periods. Estimating evolutionary rates at different time scales allows to grasp the pace of evolution. In our review, this comparison is possible for SLA, for which evolutionary rates have been estimated in *Q. faginea* and *Q. petraea*. The rates differ by two to three orders of magnitude, highlighting a faster rate of evolution when estimated over short rather than long time scales. The differences in rates are such that they can hardly be attributed to the fact that the estimates were obtained from two different species. These results suggest that estimating evolutionary rates from historical ancestral-derived approaches, can understimate the contemporary evolutionary responses of populations.

**Table 1.**
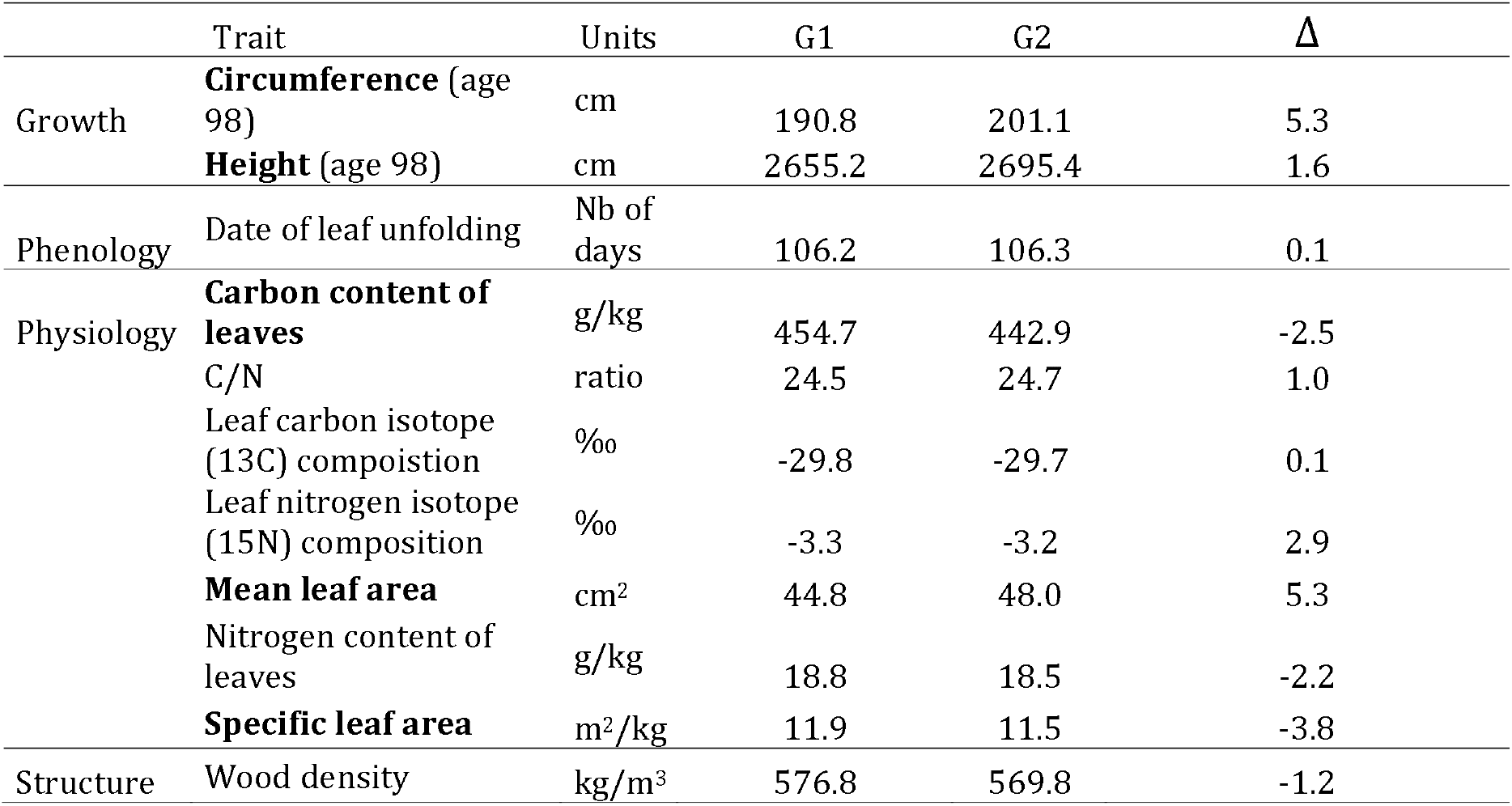
Temporal genetic changes (Δ = (G2-G1)*100/G1) of adaptive traits across two generations (G1 and G2) in *Q. petraea [107]*. Traits with significant Δ are in bold face.

| | Trait | Units | G1 | G2 | $\Delta$ |
| --- | --- | --- | --- | --- | --- |
| Growth | **Circumference** (age 98) | cm | 190.8 | 201.1 | 5.3 |
|  | **Height** (age 98) | cm | 2655.2 | 2695.4 | 1.6 |
| Phenology | Date of leaf unfolding | Nb of days | 106.2 | 106.3 | 0.1 |
| Physiology | **Carbon content of leaves** | g/kg | 454.7 | 442.9 | -2.5 |
|  | C/N | ratio | 24.5 | 24.7 | 1.0 |
|  | Leaf carbon isotope (13C) composition | ‰ | -29.8 | -29.7 | 0.1 |
|  | Leaf nitrogen isotope (15N) composition | ‰ | -3.3 | -3.2 | 2.9 |
|  | **Mean leaf area** | cm<sup>2</sup> | 44.8 | 48.0 | 5.3 |
|  | Nitrogen content of leaves | g/kg | 18.8 | 18.5 | -2.2 |
|  | **Specific leaf area** | m<sup>2</sup>/kg | 11.9 | 11.5 | -3.8 |
| Structure | Wood density | kg/m<sup>3</sup> | 576.8 | 569.8 | -1.2 |

### 4.4 Comparison of allochronic and synchronic approaches

Common garden experiments typically compare extant populations that have diverged from a common ancestral population and differentiated under contrasting environmental conditions. Hence, genetic differentiation results from both within-population evolution and evolutionary interactions among populations, notably gene flow (Fig. 3). Differentiation therefore reflects the accumulation of temporal genetic changes within each population along distinct evolutionary trajectories [20].

Intuitively, if these trajectories are shaped by the same environmental drivers and proceed in the same direction, differentiation between populations should be oriented in the same direction as evolution within populations. Consequently, differentiation and evolution are expected to be oriented in the same direction when the direction of evolutionary change is shared among populations. Conversely, if the sign of differentiation differs from that of evolution, this would suggest either contrasting evolutionary trajectories among populations or temporal fluctuations in evolutionary responses that vary across populations.

Here, in the case of *Quercus petraea*, we compare directional trends of evolution and differentiation assessed using allochronic and synchronic approaches, knowing that the time periods considered differ between the two approaches (Table 2). *Quercus petraea* is the only species for which data were available for the same traits in both allochronic and synchronic studies. We restricted the comparison to a qualitative comparison of the temperature driven trends (sign of differentiation and evolution) rather than a quantitative comparison of rates as the number of generations for assessing rates of differentiation or evolution is unknown in our studies. Temperature-driven trends of differentiation were considered positive when population mean values in common garden experiments were positively correlated with temperature at the populations’ sites of origin, and negative when the correlation was negative. Temperature-driven trends of evolution were considered positive when temporal genetic changes during warming periods were positive, and negative when changes were negative during warming periods (Table 2).

**Table 2.** Direction of within population evolutionary changes and among-population differentiation in *Quercus petraea*. The table compares the direction of temperature-associated responses for different functional traits using three approaches: a) retrospective monitoring, b) longitudinal monitoring, and c) common garden experiments. In the retrospective monitoring study, the direction of evolutionary change was assessed from temporal genetic changes associated with warming during the early Anthropocene [18]. In the longitudinal monitoring study, the direction of evolutionary change was assessed by comparing genetic values between two successive generations [107]. In common garden studies, the direction of population differentiation was assessed from the sign of the correlation between population mean trait values and mean temperature at the population sites of origin [30, 54]. Symbols indicate increasing trends (+), decreasing trends (-), no significant change or association (0), or traits not assessed (NA).

|  | Evolutionary change a) | Evolutionary change b) | Differentiation c) |
| --- | --- | --- | --- |
| Experiments | Retrospective monitoring | Longitudinal monitoring | Common gardens |
| Temporality | Early Anthropocene | Contemporary period | Holocene |
| **Growth** |  |  |  |
| Height | + | + | + |
| Diameter | + | + | + |
| **Phenology** |  |  |  |
| Date of leaf unfolding | + | 0 | - |
| Date of leaf senescence | + | 0 | 0 |
| Length of growing season | 0 | 0 | NA |
| Marcescence | - | 0 | NA |
| **Leaf morphology** |  |  |  |
| Mean leaf area (MLA) | 0 | + | 0 |
| Specific leaf area (SLA) | - | - | - |
| **Biochemistry** |  |  |  |
| Carbon content of leaves (C) | 0 | - | 0 |
| Nitrogen content of leaves (N) | 0 | 0 | 0 |
| Carbon/Nitrogen ratio (C/N) | 0 | 0 | 0 |
| Leaf Nitrogen isotopic | + | 0 | 0 |
| ($\delta^{15}\text{N}$) composition | | | |
| Leaf Carbon isotopic ($\delta^{13}\text{C}$) composition | **0** | **0** | **0** |

Sessile oak populations originated from southern European refugial areas after the last glaciation, subsequently migrated northward, and established under different climatic conditions, all of which experienced warming during the Holocene (Fig. 3). Thus, differentiation observed in common gardens reflects divergence accumulated throughout the Holocene, whereas evolutionary change assessed in the retrospective and longitudinal monitoring covers narrower time frames corresponding either to the end of the Little Ice Age and the onset of the Anthropocene (retrospective monitoring) or to the contemporary period (longitudinal monitoring). Striking similarities emerge between evolutionary changes and differentiation along both temporal and spatial temperature gradients (Table 2). Population differentiation of adaptive traits responding to temperature is oriented in the same direction as temporal genetic changes occurring within populations in response to warming. As observed in common gardens, growth, phenology, and leaf morphology traits exhibit clear evolutionary changes while most physiological traits, which are usually more plastic, do not. Moreover, these traits also display congruent directions of evolution and differentiation in response to temperature variation, with the exception of leaf unfolding. In summary, comparisons between patterns of population differentiation, assumed to have accumulated during the progressive warming of the Holocene, and within-population evolutionary changes occurring during more recent warming periods revealed remarkably similar trends. This raises the question of whether the extensive differentiation traditionally observed in common gardens of oak species developed over much shorter time scales than the entire Holocene.

### Concluding remarks

Our results showed that patterns of climatic adaptation in oak populations are strongly dependent on the biome they occupy. In temperate forests, populations from warmer and wetter areas tend to have thicker leaves, lower specific leaf area, greater leaf life span, longer growing seasons, and higher growth potential under favorable conditions. In Mediterranean environments, where summer drought is the main selective factor, populations from drier areas show more sclerophyllous leaves, lower specific leaf area, and traits associated with greater drought tolerance, such as more negative water potentials. However, as in temperate areas, growth appears to be strongly linked to the duration of the vegetative period and winter temperatures, rather than to precipitation. Tropical seasonal oaks, in contrast, show population patterns of drought response that are opposite to those expected in Mediterranean environments. In *Quercus oleoides*, for example, populations from drier environments produce larger, thinner leaves with higher specific leaf area, associated with a drought avoidance strategy through partial leaf loss during the dry season. By contrast, populations from wetter areas maintain more sclerophyllous leaves and a greater capacity for osmotic adjustment. In this biome, where there is no period of winter stress, growth is exclusively linked to precipitation patterns. Together, these results suggest that adaptation to drought does not depend only on the intensity of the dry season, but also on its interaction with other periods of stress, especially low winter temperatures.

Our results from the comparison between synchronic and allochronic approaches show that provenance tests provide relevant information on the direction of biological evolution in response to climate warming, but underestimate the rates of evolution at contemporary time scales, which are at stake in the context of ongoing climate change. These results also lead us to question the genetic drivers of rapid evolution in current tree populations. We suspect that the high levels of standing genetic variation in tree population and efficiency of natural selection, due to large available recruitment and high selection rates foster important genetic adaptive changes across a limited number of generations [108].

Forest tree species such as oaks indeed harbor high levels of additive genetic variation, which constitutes the raw material for evolution in response to natural selection. In recent years, increasing attention has been paid to common garden experiments as a way to understand patterns of population variation and potential responses to climate change, for example through space-for-time substitution approaches [4, 14]. We call for similar research efforts for experiments aiming at estimating within population genetic variation of adaptive traits, as progeny trials or *in situ* reconstruction of progenies [109].

To illustrate the evolutionary potential provided by intrapopulation variation, we show results from a common garden experiment in which six populations and a set of maternal families within each population were tested (Fig. 6, see Ramírez-Valiente et al. [110] for further details). Here, we use SLA as an example, a trait that, as shown in both synchronic and allochronic studies in previous sections, is closely linked to climate adaptation. The results show that variation among mean values of open pollinated families can be even greater than differences among population means, despite the populations being separated by hundreds of kilometers and originating from highly contrasting climates (Fig. 6). In other words, each population maintains sufficient genetic variation to ultimately generate adaptive differentiation across the distribution range of the species.

**Fig 6.**
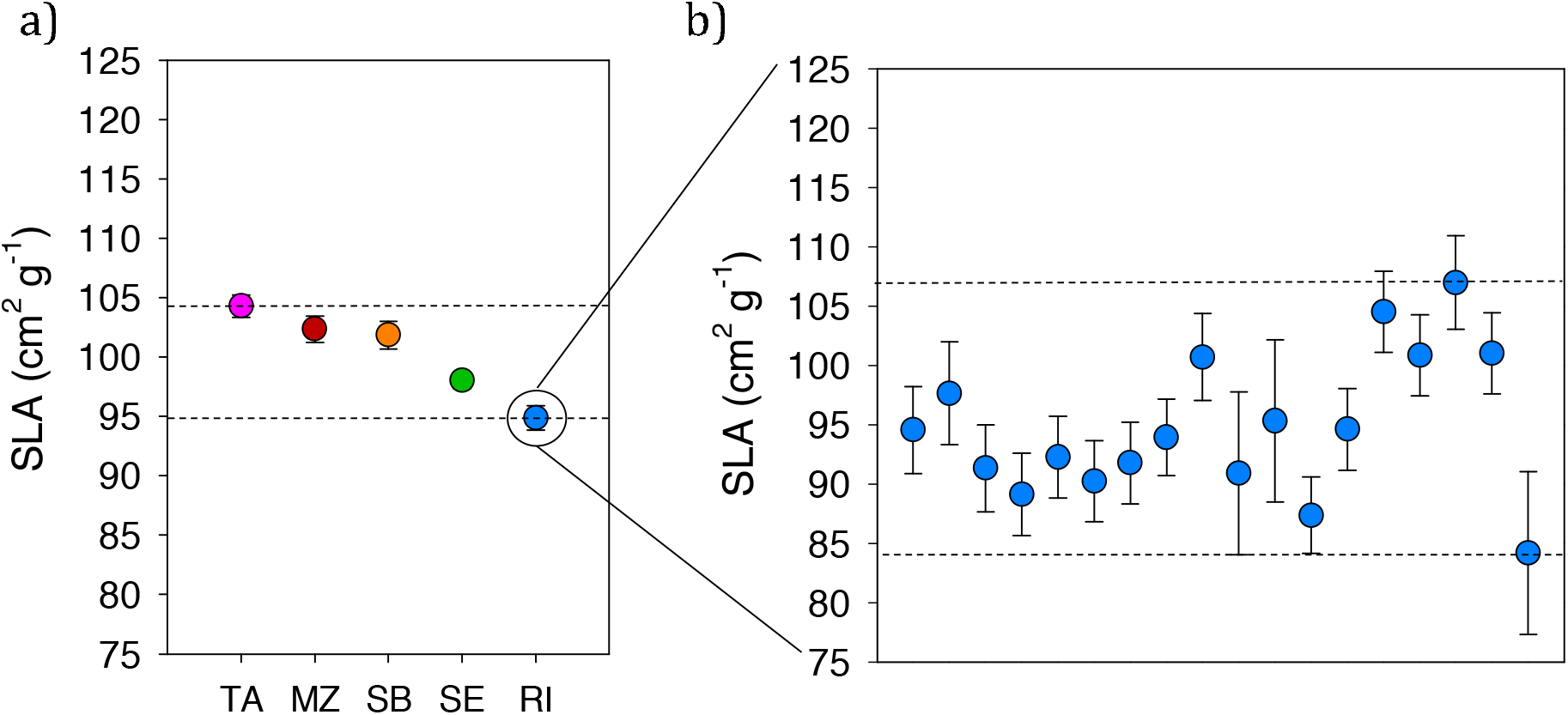
Comparison of between and within population genetic variation of SLA in *Q. oleoides*. a) Mean SLA values and standard errors for six populations of *Q. oleoides* established in a greenhouse common garden. Populations are separated by hundreds of kilometers and are characterized by differences in the duration and intensity of the dry season, ranging from approximately 2 months in RI, (Rincón de la Vieja, Costa Rica), to 6 months for TA, (Las Tablas, Honduras), see Ramírez-Valiente & Cavender-Bares [28] for further details. b) Mean SLA values for each open-pollinated family established in the greenhouse common garden within the RI population. The dotted lines in a and b indicate the range of trait variation at the population level and among open-pollinated families, respectively.

Significant within population genetic variation has also been reported for many traits in several oak species [64, 107, 111, 112]. Interestingly, while within genetic variation was traditionally estimated in designed experiments as progeny trials, which are time, resource and labor demanding, *in natura* assessments can now be implemented by reconstructing genetic relatedness directly in natural stands [109]. Such developments should facilitate the monitoring of within genetic variation and temporal genetic changes.

## Acknowledgments

This study was supported by grant 143618, funded by MICIU/AEI/10.13039/501100011033 and by the European Union NextGenerationEU/PRTR awarded to JARV. This research was also supported by the EU through an advanced ERC grant (project TREEPEACE (FP7-339728)).

## Statements and Declarations

Competing interests The authors declare no competing interests.

## Data Availability

The phenotypic data extracted from common garden studies testing population differences that were used for the meta-analysis are openly available at: https://doi.org/10.5281/zenodo.21621121.

