## Supplementary Material for "Climate adaptation across space and time: lessons from oak populations"

Table S1. Alternative demographic models tested using fastsimcoal2 for three pairs of populations of *Quercus faginea*. Models include scenarios of divergence in strict isolation (SI) and isolation-with-migration (IM, with either symmetric or asymmetric gene flow). The best-supported scenario highlighted in bold. The number of polymorphic SNPs used to calculate the site frequency spectrum for each pair of populations is indicated in parentheses.

| Model | Gene flow | lnL | *k* | AIC | ΔAIC | ω*_i_* |
| --- | --- | --- | --- | --- | --- | --- |
| (a) VAL-SAL (4987 SNPS) | | |  |  |  |  |
| SI Sy | – | -5880.24 | 2 | 11,764.48 | 164.97 | 0.00 |
| IM | Symmetric | -5797.19 | 3 | 11,600.38 | 0.87 | 0.39 |
| **IM** | **Asymmetric** | **-5795.76** | **4** | **11,599.51** | **0.00** | **0.61** |
| (b) VAL-SOT (6218 SNPS) | | |  |  |  |  |
| SI Symmetric | – | -7390.85 | 2 | 14,785.70 | 160.43 | 0.00 |
| IM | Symmetric | -7312.97 | 3 | 14,631.93 | 6.66 | 0.03 |
| **IM** | **Asymmetric** | **-7308.63** | **4** | **14,625.27** | **0.00** | **0.97** |
| (c) BOS-PEN (992 SNPS) | | |  |  |  |  |
| SI Symmetric | – | -1064.34 | 2 | 2,132.68 | 14.61 | 0.00 |
| IM | Symmetric | -1058.69 | 3 | 2,123.38 | 5.31 | 0.07 |
| **IM** | **Asymmetric** | **-1055.04** | **4** | **2,118.08** | **0.00** | **0.93** |

lnL, maximum likelihood value of the model; *k*, number of parameters in the model; AIC, Akaike’s information criterion value; ∆AIC, difference in AIC value from that of the strongest model; ω*_i_*, AIC weight.

Table S2. Parameters inferred from coalescent simulations with fastsimcoal2 under the most supported scenario of divergence – isolation-with-migration and asymmetric gene flow – for three pairs of populations of *Quercus faginea*. Table shows point estimates and lower and upper 95% confidence intervals for each parameter, which include the mutation-scaled ancestral (*θ*_ANC_) and contemporary (*θ*_VAL_, *θ*_SAL_, *θ*_SOT_, *θ*_BOS_, and *θ*_PEN_) effective population sizes, the timing of divergence (*T*_DIV_), and migration rates per generation (*m*). Estimates of time are given in number of generations. Contemporary effective population sizes were calculated from the levels of nucleotide diversity (*π*) and fixed in fastsimcoal2 analyses to enable the estimation of all other parameters (see Materials and Methods for further details).

| Parameter | Point estimate | Lower bound | Upper bound |
| --- | --- | --- | --- |
| (a) VAL-SAL | |  |  |
| *θ*_ANC_ | 1093 | 847 | 1213 |
| *θ*_VAL_ | 5070 | – | – |
| *θ*_SAL_ | 5070 | – | – |
| *T*_DIV_ | 1139 | 956 | 1354 |
| *m*_VAL🡪 SAL_ | 5.19×10^-03^ | 3.71×10^-03^ | 7.93×10^-03^ |
| *m*_VAL🡨 SAL_ | 1.44×10^-03^ | 4.43×10^-06^ | 2.47×10^-03^ |
| (b) VAL-SOT | |  |  |
| *θ*_ANC_ | 1241 | 980 | 1343 |
| *θ*_VAL_ | 5070 | – | – |
| *θ*_SOT_ | 5070 | – | – |
| *T*_DIV_ | 1257 | 1079 | 1562 |
| *m*_VAL🡪 SOT_ | 4.21×10^-03^ | 3.21×10^-03^ | 5.30×10^-03^ |
| *m*_VAL🡨 SOT_ | 8.12×10^-04^ | 3.03×10^-05^ | 1.51×10^-03^ |
| (c) BOS-PEN | |  |  |
| *θ*_ANC_ | 1062 | 209 | 1449 |
| *θ*_BOS_ | 5070 | – | – |
| *θ*_PEN_ | 4225 | – | – |
| *T*_DIV_ | 1730 | 1111 | 3182 |
| *m*_BOS🡪 PEN_ | 3.46×10^-09^ | 3.12×10^-11^ | 4.32×10^-04^ |
| *m*_BOS🡨 PEN_ | 1.97×10^-03^ | 1.29×10^-03^ | 2.66×10^-03^ |
